# Structural characterization of a pathogenic antibody underlying vaccine-induced immune thrombotic thrombocytopenia (VITT)

**DOI:** 10.1101/2023.05.28.542636

**Authors:** Son N. Nguyen, Si-Hung Le, Daniil G. Ivanov, Nikola Ivetic, Ishac Nazy, Igor A. Kaltashov

## Abstract

Vaccine-induced immune thrombotic thrombocytopenia (VITT) is a rare but extremely dangerous side effect that has been reported for several adenoviral (Ad)-vectored COVID-19 vaccines. VITT pathology had been linked to production of antibodies that recognize platelet factor 4 (PF4), an endogenous chemokine. In this work we characterize anti-PF4 antibodies obtained from a VITT patient’s blood. Intact-mass MS measurements indicate that a significant fraction of this ensemble is comprised of antibodies representing a limited number of clones. MS analysis of large antibody fragments (the light chain, as well as the Fc/2 and Fd fragments of the heavy chain) confirms the monoclonal nature of this component of the anti-PF4 antibodies repertoire, and reveals the presence of a fully mature complex biantennary N-glycan within its Fd segment. Peptide mapping using two complementary proteases and LC-MS/MS analysis were used to determine the amino acid sequence of the entire light chain and over 98% of the heavy chain (excluding a short N-terminal segment). The sequence analysis allows the monoclonal antibody to be assigned to IgG2 subclass and verify that the light chain belongs to the λ-type. Incorporation of enzymatic de-*N*-glycosylation into the peptide mapping routine allows the *N*-glycan in the Fab region of the antibody to be localized to the framework 3 region of the V_H_ domain. This novel *N*-glycosylation site (absent in the germline sequence) is a result of a single mutation giving rise to an NDT motif in the antibody sequence. Peptide mapping also provides a wealth of information on lower-abundance proteolytic fragments derived from the polyclonal component of the anti-PF4 antibody ensemble, revealing the presence of all four subclasses (IgG1 through IgG4) and both types of the light chain (λ and κ). The structural information reported in this work will be indispensable for understanding the molecular mechanism of VITT pathogenesis.

## Introduction

One of the unintended consequences of the massive vaccination campaign during the recent COVID-19 pandemic was the discovery of a rare but extremely dangerous side effect of adenoviral (Ad) vectored vaccines, vaccine-induced immune thrombotic thrombocytopenia (VITT).^1, 2^ Although VITT incidence rates are generally very low (ranging from 3 to 16 cases per million doses for ChAdOx1, and three- to four-fold lower for human Ad-vectored vaccines^3^), some studies reported notably higher numbers (e.g., 5 cases per 132,686 vaccine doses in Norway^4^). Furthermore, VITT is associated with very high mortality rates,^1^ and it certainly became one of the factors contributing to the vaccine hesitancy phenomenon and undermining the COVID-19 vaccination campaign globally.^5^ Lastly, the close association of VITT with a specific delivery vector naturally invites the question of whether this side effect should be expected for other Ad-vectored vaccines (both existing and those that are still at the development stage), an alarming prospect given the growing popularity of this platform.^6^

Despite having several distinct features, such as thromboses at unusual sites (including cerebral venous sinus thrombosis^7^), clinical presentation of VITT is strikingly similar to another immune-mediated blood disorder, heparin-induced thrombocytopenia (HIT).^8^ At the molecular level, both VITT and HIT are associated with the emergence of antibodies recognizing a small chemokine, platelet factor 4 (PF4).^4, 9^ Since anti-PF4 antibodies play a central role in HIT pathogenesis, structural characterization of VITT-associated anti-PF4 antibodies may shed light on the etiology of this vaccination side effect, as well as offer viable treatment and prophylaxis options. The initial mapping of the VITT antibodies’ epitopes on the PF4 surface using alanine scanning revealed their significant overlap with the heparin-binding sites,^10^ leading to a suggestion that the clonality of these antibodies is significantly restricted. Subsequent work by other groups confirmed that the anti-PF4 antibodies in VITT patients are generated by a very limited number of B cells based on the mass profiling of their light chains^11^ and amino acid sequences of their CDR3 regions.^12^ The purpose of this study is complete structural characterization of a monoclonal anti-PF4 antibody extracted form a VITT patient’s blood (including not only the amino acid sequencing, but also localization of all glycans – including those outside of the canonical *N*-glycosylation site within the C_H_2 IgG domain^13^) with an overarching goal of generating a molecularly defined model of VITT pathogenesis. While in the past the antibody sequencing task was approached by gene sequencing of the B cell receptors at a single clone level,^14^ recent advances in mass spectrometry enabled de novo sequencing of endogenous antibodies at the protein level,^15^ although a range of technical issues remain to be addressed prior to this method becoming a routine tool in clinical analysis.^16^

## Materials and Methods

### Extraction and purification of anti-PF4 antibodies from a VITT patient’s serum

The plasma sample used in this study was obtained from a clinically diagnosed VITT patient referred for diagnostic testing to the McMaster Platelet Immunology Laboratory. The VITT diagnosis was based on (i) recent AstraZeneca vaccination, (ii) presence of thrombotic complications, (iii) testing serologically positive for anti-PF4 IgG antibodies using the commercially available PF4-enhanced heparin-dependent IgG/A/M-specific EIA Immucor, OD ≥ 0.45), and (iv) positive PF4-enhanced serotonin release assay, SRA (SRA ≥ 20% ^14^C-serotonin release). This study was approved by the Hamilton Integrated Research Ethics Board (HiREB). Anti-PF4 antibodies were purified from the patient plasma using the following protocol. Plasma was heat-treated at 56 ^O^C for 30 min. and centrifuged at 21,000 g for 10 min. to separate the sample supernatant. The isolated supernatant was diluted 4-fold with phosphate buffered saline (PBS) and loaded onto a gravity flow column containing protein G sepharose beads (Protein G Sepharose 4 Fast Flow cat 17061801, Cytiva, Marlborough USA). The beads were washed with PBS until the optical density of the flowthrough was observed to be < 0.09 at 280 nm. The retained antibodies were then eluted off the column using glycine (0.1 M, pH 2.8) and immediately neutralized using tris (2 M, pH 8.4). The eluted total IgG was buffer exchanged to PBS and concentrated to 7 mg/mL using 50 kDa centrifugal filter units (Ultracel-50, cat UFC905024, Amicon, Tullagreen IRL). The purified IgG fraction was incubated with biotinylated PF4 (0.1 mg/mL final concentration, generated inhouse as previously described,^17^ for 2 hrs at room temperature under gentle rocking conditions. Washed streptavidin sepharose beads (Streptavidin Sepharose High Performance, cat 17511301 Cytiva, Marlborough USA) were added to the PF4/antibody mixture (at a ratio of 1.5 mL 50 % slurry to 2 mL antibody solution) and left to incubate overnight on a rocker at 4 ^O^C. Beads were then washed with PBS till the optical density was < 0.09 at 280 nm. The bound antibody was then eluted off the beads as described before using glycine and tris. The eluted IgG was then mixed with 2 M NaCl at a 1:10 volume ratio and incubated for 15 min. on a rocker. The sample was then centrifuged at 300 g, 1 minute and filtered through a 0.2 µm filter to remove beads before being buffer exchanged to PBS using a 50 kDa centrifugal filter. The final sample was then tested in the PF4-enhanced SRA to confirm reactivity.

### Mass spectrometry

Intact-mass measurements of the VITT patient-derived anti-PF4 antibodies were carried out with a Synapt G2 HDMS (Waters Corp., Milford, MA) hybrid quadrupole/time-of-flight mass spectrometer equipped with a NanoLockSpray ion source. The anti-PF4 antibodies PBS solution was buffer-exchanged to 150 mM Ammonium Acetate, pH 6.8 and loaded into the gold-coated borosilicate capillaries. The following nanoESI source parameters were used to maximize ion desolvation in the ESI interface and minimize the signal generated by non-covalent adduct ions: source temperature, 30 ^O^C; capillary-voltage, 1.5 kV; sampling cone, 200 V; and extraction cone, 4 V. Charge state assignment was carried out using limited charge reduction,^18^ and these values were used as input for UniDEC^19^ to obtain the antibody mass distributions by deconvolution.

Mass analyses of large antibody fragments (the Fc/2 and Fd segments of the heavy chain, and the light chain) were carried out with a SolariX 7 (Bruker Daltonics, Billerica, MA) Fourier transform ion cyclotron resonance (FT ICR) MS in the LC/MS mode. The large fragments were generated by digesting the antibody sample with IdeZ (ThermoFisher, Waltham, MA) for 1 hr at 37 ^O^C using 1 µL of the enzyme (8 units/µL) for 5 µg of the substrate, followed by disulfide reduction with 0.3 M DTT (Millipore Sigma, St. Louis, MO) in Glycobuffer 2×10 solution maintained at 55 ^O^C for 1 hr in the dark. Antibody de-*N*-glycosylation was carried out by incubating the sample with PNGase F (New England Biolabs, Ipswich, MA) for 24 hrs at 37 ^O^C (enzyme:substrate ratio *ca*. 100 units:1 µg) prior to the IdeZ digestion step. The fragments were separated on a 4.6×100 mm, 3.5 µm pore size AdvanceBio RP-mAb column (Agilent Technologies, Santa Clara, CA) at room temperature, and the eluate was directly injected into the ESI source of the FT ICR MS at a 300 µL/min flow rate. The ESI interface temperature was maintained at 250 ^O^C to maximize ion desolvation. The UniDEC^19^ algorithm was used for deconvolution of the mass spectra.

Amino acid sequencing and glycan localization were carried out by digesting the antibody samples with trypsin or chymotrypsin (New England Biolabs, Ipswich, MA) followed by LC-MS/MS analysis of proteolytic fragments with an Orbitrap Fusion (Thermo, San Jose, CA) LC/MS system equipped with an on-line Easy-nLC 1000 chromatographic system and a NanoSpray Flex ion source. Fragmentation of peptide ions was carried out using HCD. The initial data processing was carried out with a PEAKS Studio Xpro (Bioinformatics Solutions, Waterloo, ON) software suit, followed by manual inspection of fragment ion assignment in all relevant mass spectra.

## Results and Discussion

### Intact-mass analysis of anti-PF4 antibodies in the clinical sample reveals the presence of an abundant monoclonal antibody with an anomalously high mass

The anti-PF4 IgG molecules extracted from a VITT patient’s blood were subjected to intact-mass analysis under near-native conditions in solution and aggressive desolvation in the gas phase. This allowed the spectral crowding (typically observed in the low *m/z* regions of mass spectra of highly heterogeneous proteins acquired under denaturing conditions^20^) to be avoided, while minimizing the presence of signal corresponding to non-covalent adduct ions.^21^ Even under these conditions the anti-PF4 antibody mass spectrum displays a highly convoluted ionic signal (**Figure 1A**), and its processing even with the most robust deconvolution algorithms without applying any additional filters/restrictions yields ambiguous results. To resolve this ambiguity, the limited charge reduction technique^22^ was applied to multiple ionic populations selected within narrow *m/z* windows, revealing the presence of three distinct components (**Figure 1B**). The charge state range (*z* = +23 through +30) provided by the limited charge reduction measurements was used as an additional filter to restrict the search space for the UniDEC deconvolution, yielding a distribution spanning a 145-153 kDa mass range (inset in **Figure 1A**). The deconvoluted mass distribution has a clearly defined bimodal character, with the lower part of the distribution having an appearance that is typical of polyclonal antibodies (in fact, it juxtaposes well with a deconvoluted mass distribution of anti-PF4 antibodies extracted from a HIT patient’s blood, which are known to be polyclonal^23^ – see the purple trace in the **Figure 1A** inset). In contrast to this part of the distribution, the higher-mass component of the signal has a well-defined shape typical of monoclonal antibodies,^24^ although the level of resolution and mass accuracy is not sufficient to conclude with certainty whether it represents partially resolved glycoforms of a monoclonal antibody or a small number of oligoclonal antibodies. Another notable feature of the high-mass component of the anti-PF4 antibody signal is the anomalously high mass of the corresponding IgG molecules.

**Figure 1.**
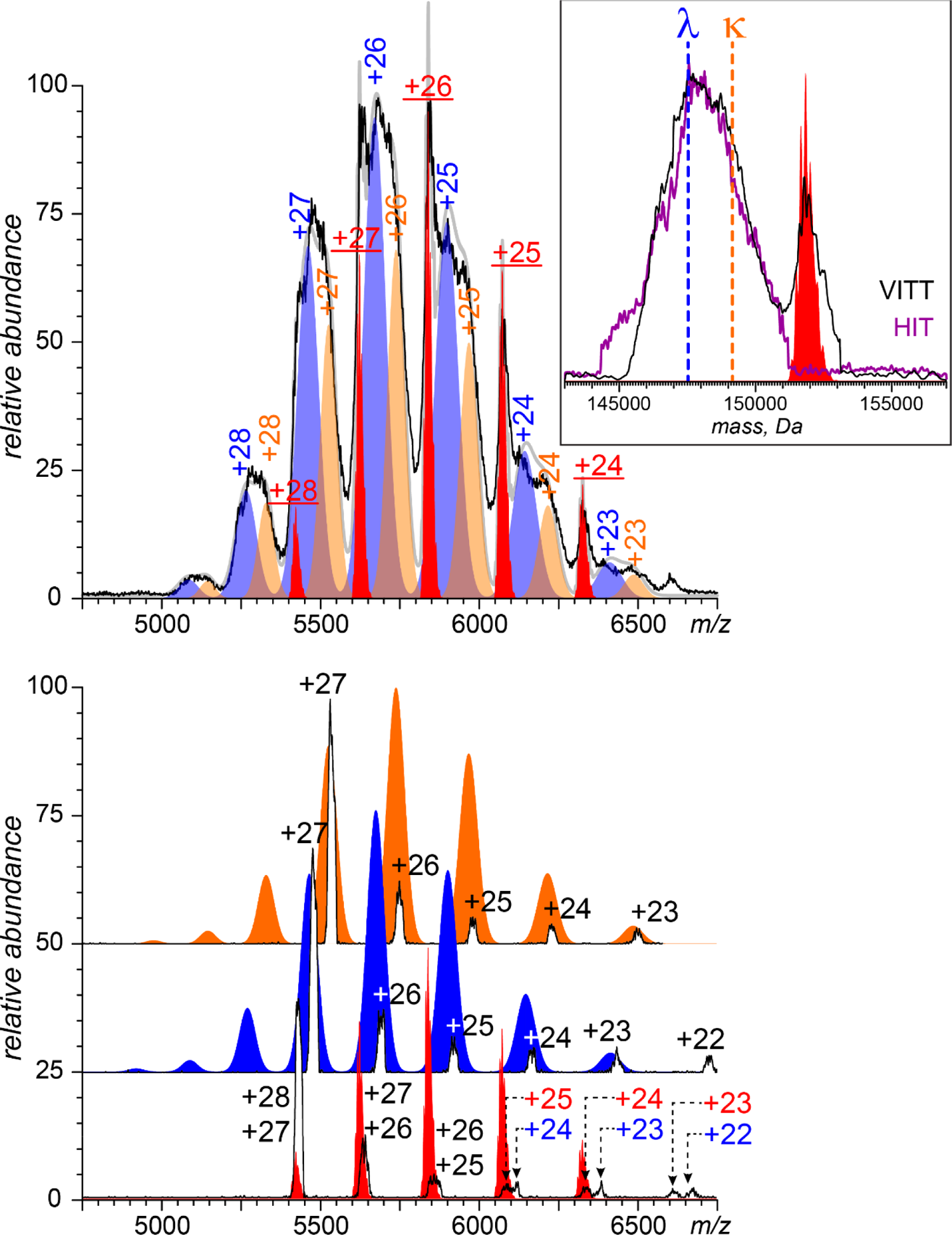
Intact mass analysis of anti-PF4 IgG antibodies extracted from the blood of a VITT patient. (**A**) and limited charge reduction measurements (**B**) carried out to facilitate the ionic charge state assignments. The inset shows deconvoluted mass spectra of the anti-PF4 antibodies extracted from the VITT patient’s blood (black trace) and HIT-associated anti-PF4 antibodies as a reference (purple). The color-filled curves show the results of data fitting with three components: red, monoclonal (obtained by convolution of mass distributions of the its light chain and Fc/2 and Fd fragments of the heavy chain, see Figure 2); blue, polyclonal λ-component; and orange, polyclonal κ-component.

Indeed, while the appearance of the lower-mass component of the antibody signal is consistent with the masses expected for the two most common subclasses (IgG1 and IgG2) carrying light chains of both λ- and κ-types^25^ and a single biantennary glycan of a complex type per each heavy chain (see the data fitting in **Figure 1A**), the notably higher mass of the component representing the antibodies with restricted clonality must invoke different interpretations. One possibility is that this component is represented by an IgG molecule of a less common subclass - IgG3 – which features an elongated hinge region with multiple O-glycans.^26^ Another possible explanation of the mass anomaly is the presence of additional *N*-glycans within these molecules outside of the canonical N-glycosylation site within the C_H_2 domain.^27^

### Mass profiling of large antibody fragments reveals the presence of a fully mature biantennary N-glycan of the complex type within the Fab segment of the VITT patient-derived anti-PF4 mAb

IdeZ digestion of the anti-PF4 antibody sample followed by disulfide reduction and LC-MS analysis of the resulting fragments yielded mass distributions of the intact light chain, as well as the Fd and Fc/2 fragments of the heavy chain (the former comprising the V_H_, C_H_1 domains and the hinge region, and the latter comprising the C_H_2 and C_H_3 domains, see **Figure 2**). Interestingly, fragmenting the entire anti-PF4 antibody ensemble to the level of these segments resulted in a dramatic reduction of the relative abundance of the polyclonal component of the signal. In fact, the mass distribution of the reduced light chain is dominated by a single peak at 22,678±2 Da, a mass consistent with the λ-type.^25^ Unlike the light chain, mass distributions of both Fd and Fc/2 fragments of the antibody reveal some heterogeneity (see the blue traces in the corresponding panels of **Figure 2**). While it might be explained away by invoking the notion of a limited number of clones contributing to the overall signal, we note that the spacing patterns between the adjacent peaks in these deconvoluted mass spectra are consistent with the notion of limited heterogeneity caused by the presence of glycans. This pattern is fully expected for the Fc/2 segment (where the “internal” placement of the *N*-glycan within the C_H_2 domain frequently results in incomplete – and heterogeneous – glycosylation due to the steric constraints imposed upon the relevant enzymes by the antibody architecture). The two abundant peaks in the Fd mass distribution are also separated by a mass increment corresponding to a HexNAc residue, but it is not clear whether this pattern is indicative of the presence of a non-canonical *N*-glycan chain within this segment of the heavy chain or reveals the O-glycosylation within the hinge region, typical of IgG3 antibodies.^26^ Repeating these measurements using stressed anti-PF4 antibodies obtained in the course of plasmapheresis (red traces in **Figure 2**) had no effect on the measured mass distribution of the light chain, while revealing a few additional glycoforms within the Fd and Fc/2 segments, consistent with the limited degradation of the carbohydrate chains (but insufficient to determine the glycan type within its Fd segment).

**Figure 2.**
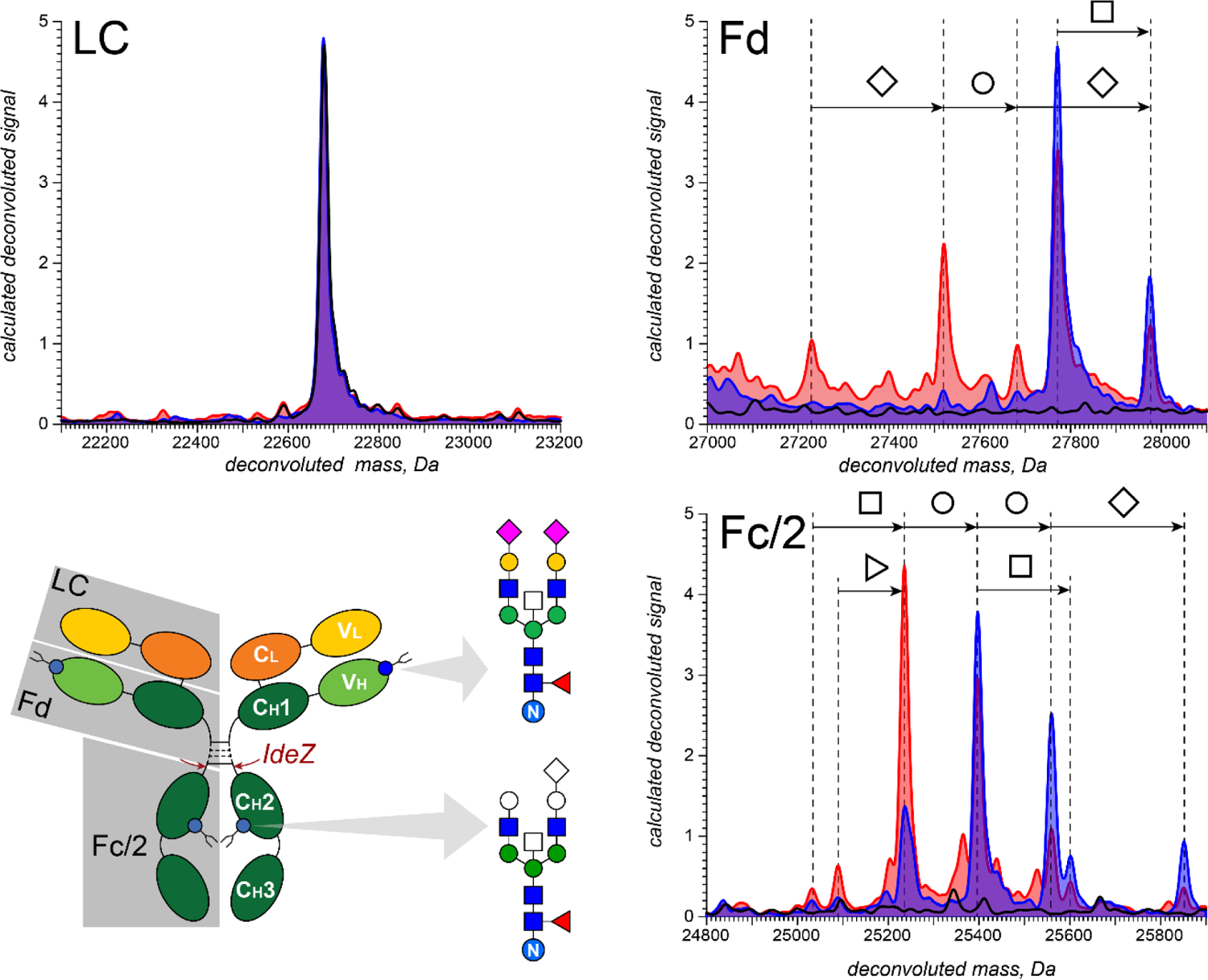
Deconvoluted mass spectra of the light chain, Fd segment of the heavy chain and Fc/2 segment of the heavy chain of VITT mAb derived by IdeZ digestion and disulfide reduction from intact (blue-filed curves) and stressed (red-filled curves) of the anti-PF4 antibodies extracted from the VITT-patient’s blood (the stressed sample was obtained from the plasmapheresis procedure). The black traces show the effect of de-N-glycosylation on the mass spectra of these segments. The schematic representation of the two N-glycans (***bottom left***) shows the color-filled base structures (corresponding to the lowest-mass glycoform detected by MS); the higher-mass glycoforms are shown with additional colorless monosaccharide symbols.

The nature of the glycan residing within the Fd segment of the monoclonal anti-PF4 antibody was determined by de-*N*-glycosylation of the antibody with PNGase F prior to its processing with IdeZ and disulfide reduction. As expected, the PNGase treatment had no effect on the mass distribution of the light chain (black trace in **Figure 2**). It also confirmed the presence of an immature biantennary *N*-glycan of the complex type within the Fc/2 segment (whose mass distribution collapsed to a single peak at 23,792±3 Da). Importantly, the mass distribution of the Fd segment also collapsed to a single peak (25,422±3 Da), not only confirming the presence of an *N*-glycan within this segment, but also revealing the identity of the major glycoform as a fully mature biantennary *N-*glycan of the complex type (based on the close match of the mass difference between the most abundant isoform of Fd before and after PNGase F treatment – 2,350±3 Da – and the calculated mass of the Fuc_1_Hex_5_HexNAc_4_NeuAc_2_ chain of 2,350.9 Da). We also note that the second most abundant Fd glycoform is a result of an addition of a fifth HexNAc residue to the complete biantennary chain, the so-called “bisecting” GlcNAc, consistent with the previous reports of the structure of *N*-glycans found in Fab segments of IgG molecules.^13^ These measurements clearly confirm the monoclonal nature of the homogeneous high-mass component of the anti-PF4 antibodies extracted from the VITT patient’s blood sample, which will be referred to as VITT mAb in the subsequent sections. In addition, these measurements reveal the presence of a fully mature biantennary N-glycan within the Fab segment of VITT mAb, a feature that had been noted in the past to be correlated with the emergence of monoclonal antibodies in several auto-immune disorders,^28, 29^ as well as placental immune evasion.^30^ In addition, Fab glycosylation has a potential to impact a range of physical and biochemical properties of IgGs (such as stability),^31^ many of which may have important – although not completely understood – clinical implications.^13^ Importantly, convolution of the mass distributions of the large fragments of VITT mAb shown in **Figure 2** produces a mass distribution that juxtaposes with the high-mass component of the anti-PF4 VITT antibodies (**Figure 1**), confirming that the molecular entity represented by this component is indeed VITT mAb.

While the aberrant production of monoclonal antibodies is known to occur in a range of autoimmune disorders, understanding both the molecular mechanism of VITT pathogenesis and the etiology of this dangerous immune response would not be possible without detailed structural characterization of the VITT mAb. While the amino acid sequence of the complementarity defining regions (CDRs) within the V_L_ and V_H_ domains of the antibody are obviously the major determinant of its antigen affinity and specificity, other structural aspects are important as well. For example, identification of the antibody subclass (IgG1, IgG2, IgG3 or IgG4) is important vis-à-vis clarifying the specificity of its interaction with a range of relevant Fcγ receptors,^32^ most notably FcγRIIa on the platelet surface. Furthermore, identification of the *N*-glycosylation site within the Fab segment of the monoclonal antibody may catalyze the search for molecular mechanisms enabling selection of the aberrant clone producing these pathogenic IgG molecules.^33^ Since isolation of the clone was not feasible within the framework of this study, de novo sequencing of VITT mAb provides the only means to answer these questions.

### The hinge region structure determination allows the monoclonal anti-PF4 antibody subclass to be identified as IgG2 and confirms the light chain type λ

Amino acid sequencing of and *N*-glycan localization within the VITT mAb was carried out using peptide mapping and LC-MS/MS analysis of the proteolytic fragments. Trypsin and chymotrypsin were used to produce two complementary peptide maps, which facilitated the fragment peptide alignment (**Figure 3**). The PEAKS algorithm^34^ was used for the initial peptide sequence assignment, which were then verified/corrected manually (*vide infra*). Since the proteolytic fragments were derived from the entire ensemble of anti-PF4 antibodies *(i.e*., VITT mAbs were not separated from the polyclonal component prior to the proteolytic processing), the resulting peptide fragment mixtures contained a lower-abundance, but extremely diverse background representing the polyclonal anti-PF4 antibodies. Typically, these ions’ abundance fell 1-2 orders of magnitude below that of the peptide ions representing the VITT mAb, which allowed an intensity-based criterion to be used as a filter. The VITT mAb sequence assembly and verification was greatly facilitated by the availability of the germline gene sequences,^35^ which made the peptide alignment within the constant regions of the antibody relatively straightforward. Indeed, while the mass measurement of the VITT mAb light chain (**Figure 2**) suggests that it belongs to the λ-type (based on the mass distributions of the λ- and κ-light chains compiled by Barnidge et al.,^25^ it is the abundant tryptic and chymotryptic fragments derived from the C_L_ domain of the antibody (**Figure 3**) that allow its type to be unequivocally established as λ based on the sequence alignment with the corresponding gene IGLC2^35^ (see ***Supplementary Material*** for more detail).

**Figure 3.**
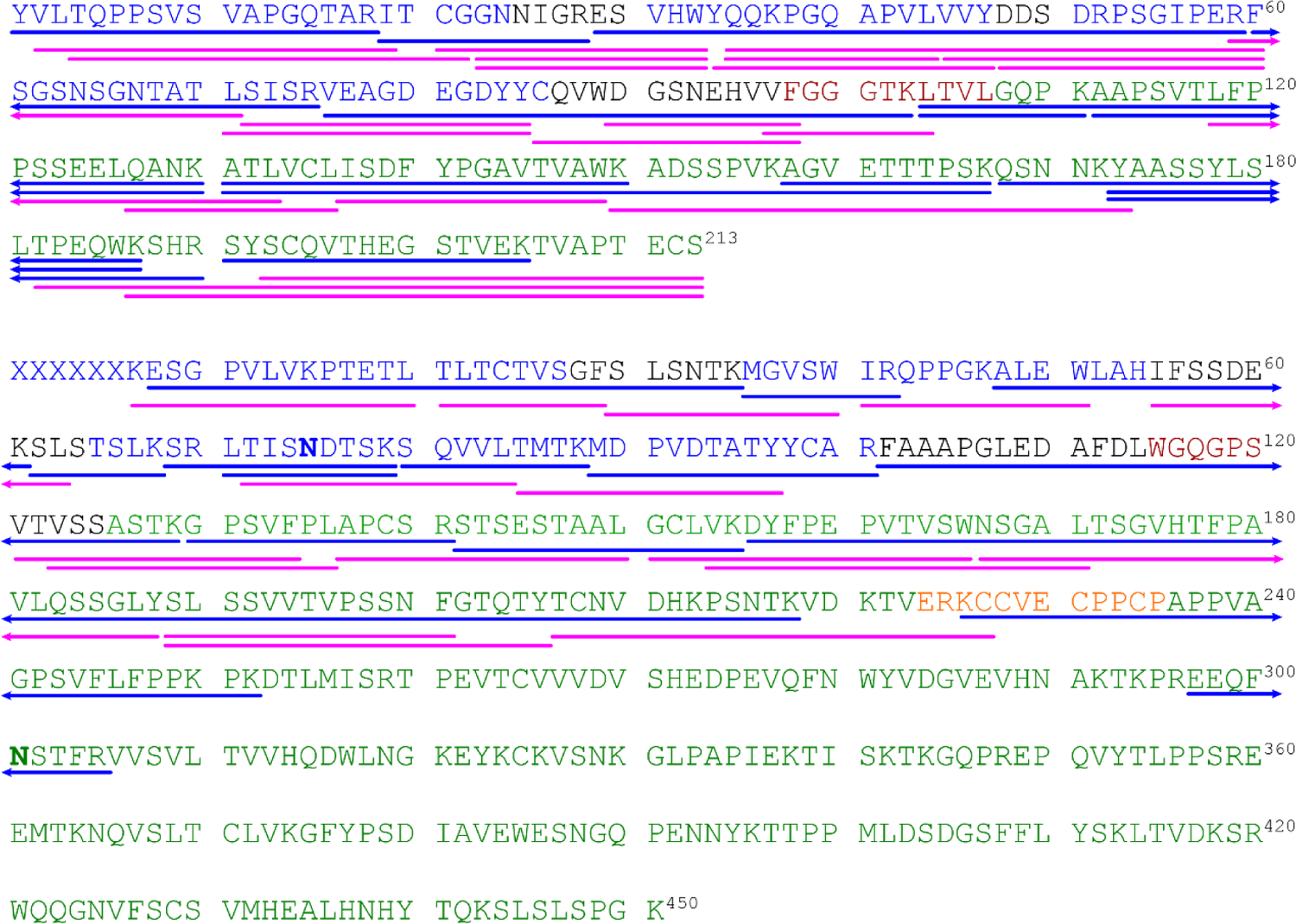
Peptide maps of light (***top***) and heavy (***bottom***) chains of the monoclonal anti-PF4 antibody extracted from the VITT patient’s plasma. CDR segments are shown in black, FRs in blue, V/C junctions in maroon, constant domains in green and the heavy chain hinge region in gold. The *N*-glycosylation sites within the V_H_ and C_H_2 domains are typed in bold. The identified tryptic and chymotryptic peptide fragments are shown with blue and magenta lines, respectively (the only proteolytic fragments shown in the C_H_2 and C_H_3 domains of the heavy chain are those used for the IgG sub-class identification).

The constant regions of the heavy chain exhibit more diversity due to the existence of four distinct sub-classes (IgG1 through IgG4), and while the most dramatic variability is exhibited by the hinge region, some differences exist within the C_H_1, C_H_2 and C_H_3 domains as well.^35^ **Figure 4** shows the fragment ion spectrum of the peptide that was tentatively identified by PEAKS as part of the C_H_1 domain, although a large number of abundant fragment ions were not automatically identified/assigned in the corresponding mass spectrum. A manual inspection of this spectrum revealed the presence of clusters of high-intensity peaks spaced by 18.011±0.001 Da, a mass difference corresponding to multiple (up to three) losses of H_2_O molecules. While such large numbers of H_2_O molecules eliminated from fragment ions are relatively rare and are usually not included in search algorithms, we note that the tentative assignment of this peptide as SLSSVVTVPSSNF provides an explanation of this phenomenon. Indeed, this peptide contains a large number of residues with the side chains terminating in hydroxyl groups, and the extent of H_2_O elimination for each fragment is clearly correlated with the number of such residues (serine and threonine) it encompasses (**Figure 4**). Even though this peptide maps to the C_H_1 domain, the two C-terminal residues of this peptide are unique to the IgG2 subclass, allowing the VITT mAb to be tentatively assigned to this subclass.

**Figure 4.**
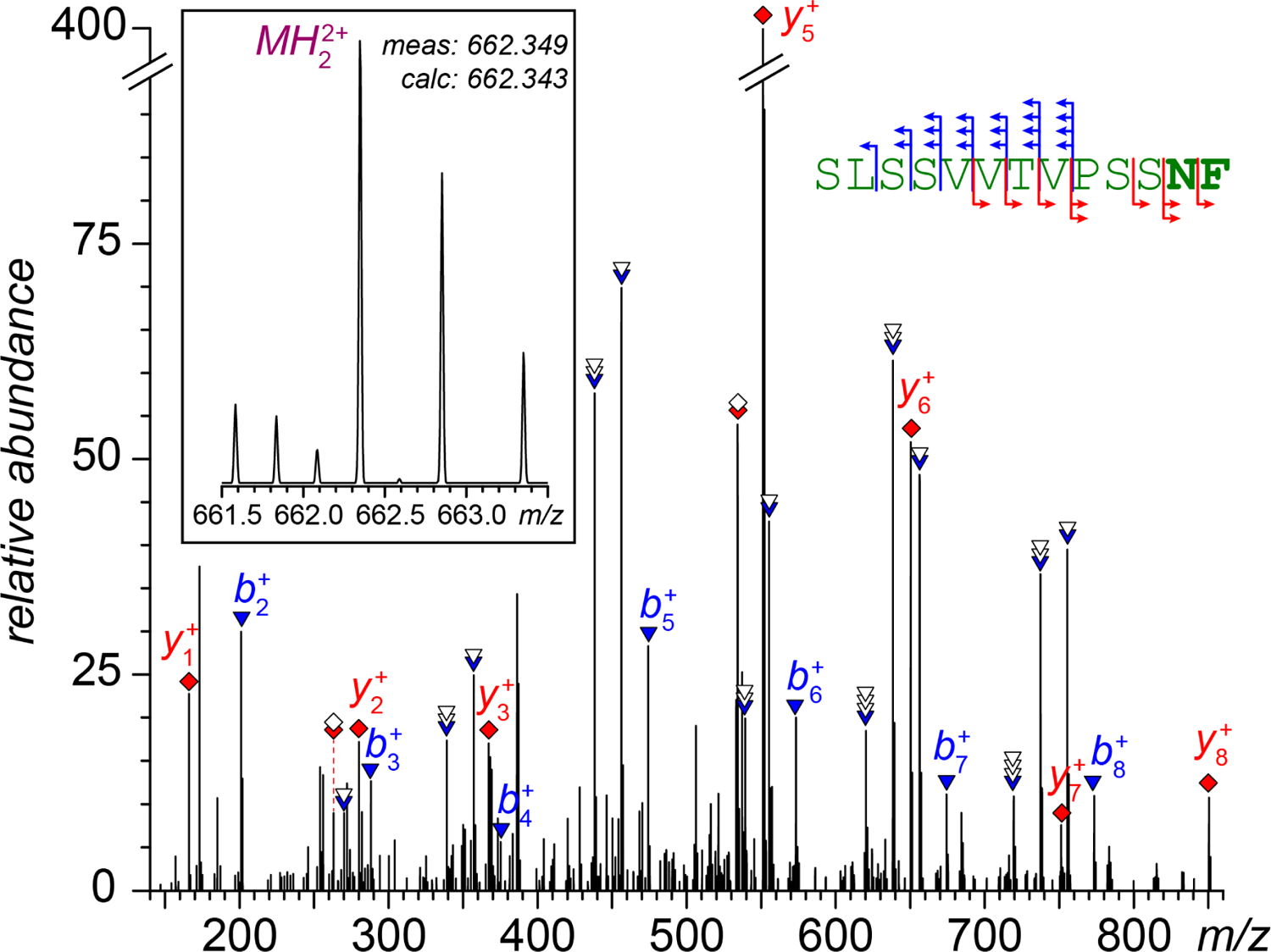
LC-MS/MS identification of a chymotryptic fragment SLSSVVTVPSSNF derived from the VITT mAb C_H_1 domain and unique to the IgG2 subclass (the two C-terminal residues for other subclasses are SL. Ionic peaks corresponding to *b*-fragments are labeled with blue triangles (overlaid white triangles represent loss of water molecules), and those corresponding to *y*-fragments are labeled with red diamonds (overlaid white diamonds represent loss of an ammonia molecule). All fragment masses match the calculated ones to 10 ppm. The inset shows the ionic signal of the precursor ion without collisional activation.

Further evidence allowing the VITT mAb to be classified as IgG2 comes from the sequence of the hinge region. For example, an abundant peptide spanning the entire hinge region and the N-terminal part of the C_H_2 domain (**Figure 5**) reveals the amino acid sequence and the high linear density of cysteine residues characteristic of this subclass. The fragment ion assignment in this case is relatively straightforward due to the paucity of hydroxyl-containing residues (and hence the absence of fragment ions generated by H_2_O elimination), and the presence of multiple proline residues (giving rise to abundant fragments). In this case the manual inspection of the fragment ion spectrum does not yield any additional features beyond those generated automatically by the search algorithm, and simply confirms all assignments.

**Figure 5.**
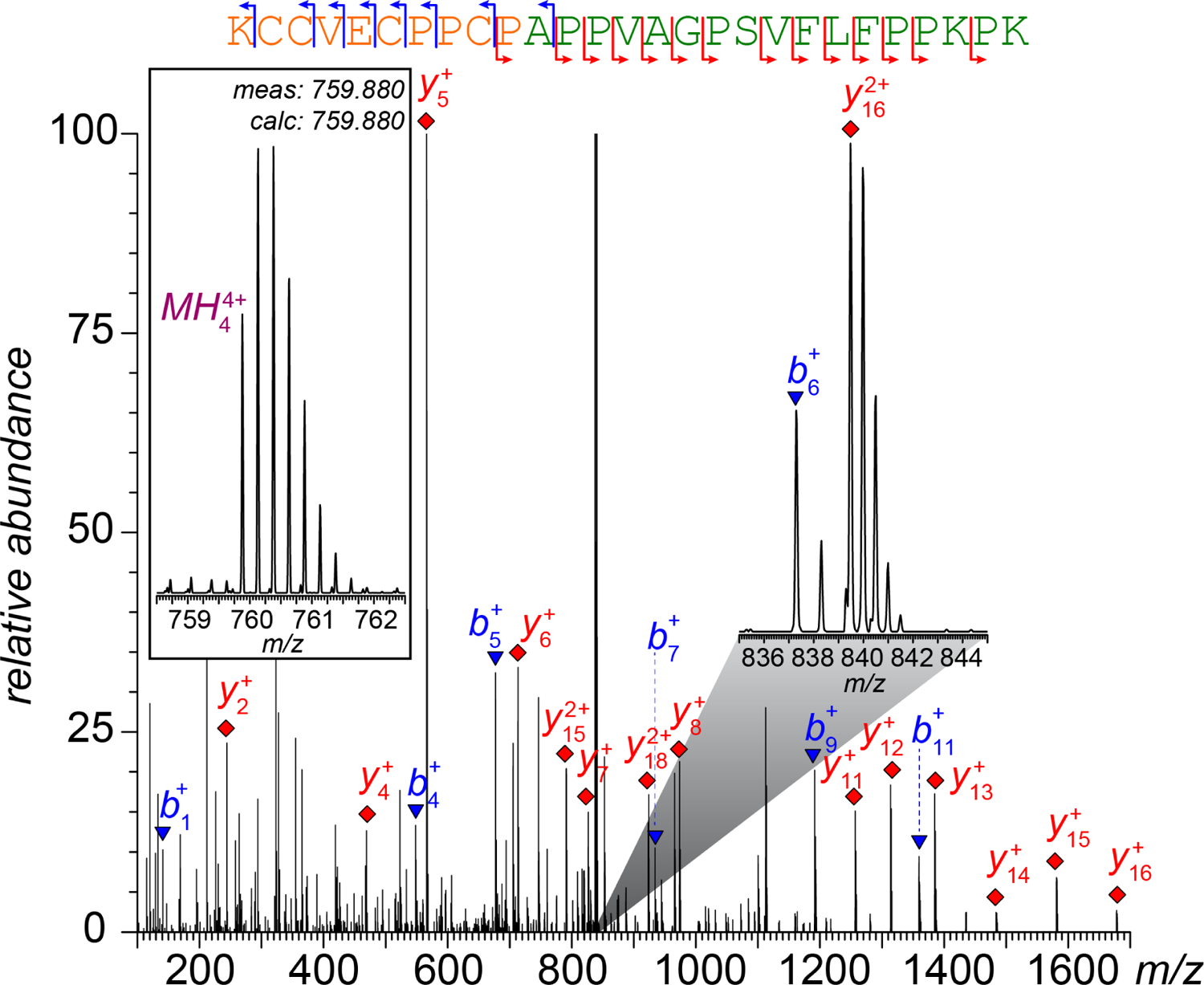
Identification of a tryptic fragment KCCVECPPCPAPPVAGPSVFLFPPKPK covering the hinge region and the N-terminal part of the VITT mAb C_H_2 domain (unique to the IgG2 subclass). Ionic peaks corresponding to *b*- and *y*-fragments are labeled with blue triangles and red diamonds, respectively. All fragment masses match the calculated ones to 10 ppm. The zoom-in 837-854 m/z region shows the detail of the partially overlapping isotopic distributions of the two most abundant fragment ions (over-scale on the main graph), and the framed inset on the left shows the ionic signal of the precursor ion without collisional activation (the sequence coloring scheme follows that shown in Figure 3).

### CDR sequencing provides structural information on the VITT mAb regions defining its antigen specificity

Amino acid sequencing of the VITT mAb variable regions (both V_L_ and V_H_) can be somewhat assisted by the availability of the germline sequences of the IGLVX (X= 1-11), IGLJX (x = 1-7), IGHVX (X=1-8) and IGHJX (X=1-6) genes and their alleles,^35^ which may provide valuable reference points for mapping the CDR regions. At the same time, one needs to be mindful of the possibility of somatic hypermutations affecting not only the CDR segments, but also the framework regions (FRs) and V/C junctions in both chains, thereby complicating the use of the gene sequence information. Because of these considerations, amino acid sequences of the CDR regions were determined using de novo sequencing approaches, followed by use of the germline gene sequences to verify CDR placement/assignment.

An example of using de novo sequencing as a means of identifying the LCDR3 (usually the most variable part within the V_L_ domain)^36^ is presented in **Figure 6A**. Despite the large size of this peptide (spanning - in addition to the entire LCDR3 – the C-terminal part of the FR3 segment and most of the V_L_/C_L_ junction region), abundant fragment ions provide a nearly-complete coverage across the entire sequence, which allows its placement to be made readily. In this case, the *y*-ions ladder covers the entirety of the peptide sequence with only two gaps, at *y*_25_ and *y*_27_ positions. The small number of *b*-ions detected/assigned automatically and verified manually do not fill these gaps, either. While the sequence ambiguity created by one of these gaps (due to the absence of *b*_4_/*y*_25_ fragments) is small and affects only the order of two residues in the sequence (GD *vs*. DG), the uncertainty created by another gap (*b*_1_/*y*_27_ fragments) provides the following four options based on the measured difference between the masses of the precursor ion and the shortest detected *b*-fragment: VE, LD, EV and DL (here we assume that L and I are identical residues, since the ion activation methods used in this work do not allow a distinction between them to be made). However, careful examination of the fragment ion peaks unassigned by the PEAKS algorithm reveals the presence of three groups of internal ions (see **Figure 6A**). The largest of these groups creates a ladder spanning the EAGDEGDYY segment at a single residue-level resolution, providing the amino acid sequence information for both gaps. Therefore, a combination of the de novo sequencing algorithms with manual data processing allows the entire amino acid sequence of this peptide to be determined.

**Figure 6.**
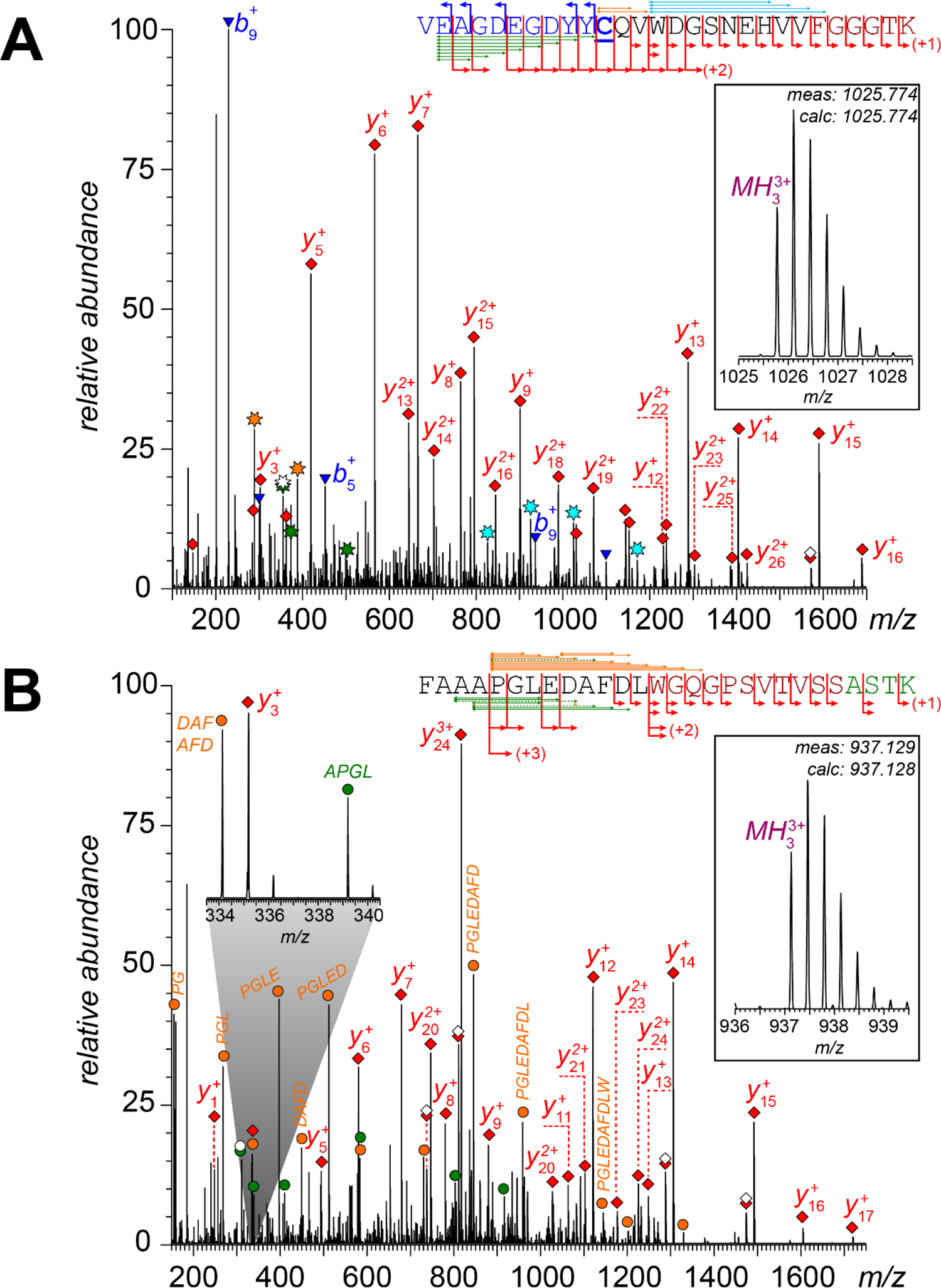
LC-MS/MS analysis of tryptic peptides covering the LCDR3 (**A**) and HCDR3 (**B**) regions of the VITT mAb (the framed insets in both panels show the ionic signal of the intact peptide ions selected as precursors for MS/MS measurements in each case). Ionic peaks corresponding to *b*- and *y*-fragments are labeled with blue triangles and red diamonds, respectively. Insertion of an open symbol indicates either H_2_O or NH_3_ loss from the corresponding fragment ion. Internal fragments are designated with gold, olive and teal symbols (placement of all internal ions within the peptide sequence is indicated with the appropriately colored double-headed arrows, with dotted lines indicating ambiguous assignments). All fragment masses match the calculated ones to 10 ppm. The zoom-in 333.5-340.5 *m/z* region in panel **B** shows the detail of partially overlapping isotopic distributions of two representative fragment ions, as well as one of the internal fragments critical for establishing the amino acid order within the four-residue long N-terminal segment). The peptide sequence coloring scheme follows that shown in Figure 3.

Interestingly, among the four deviations form a germline sequence within this antibody segment, one (A7G) falls outside of the CDR3 and is located in the FR3, where mutation frequency is generally much lower compared to CDR’s.^36^ This highlights the need to exercise caution when using gene sequences to facilitate the CDR sequencing/localization work. Nevertheless, the close match of the rest of the N-terminal sequence of this peptide to that of FR3, the presence of a cysteine residue and the match of the C-terminal segment of the peptide to the six residue-long part of the V_L_/C_L_ junction sequence, as well as the canonical 11 residue length of the CDR segment allow the latter to be confidently assigned as LCDR3.

Likewise, HCDR3 (usually the most variable segment of the entire antibody)^36^ is presented by a peptide spanning its entire length, and also including the V_H_/C_H_ junction region and a short N-terminal segment of the C_H_1 domain (**Figure 6B**). While the automatically generated sequence shows complete coverage in this case, the abundance of the *b*-fragments used to identify the first four *N*-terminal residues was determined to be extremely low upon the manual inspection of the data. As a result, these fragment ions were discarded, leaving uncertainty vis-à-vis the *N*-terminal segment composition. Another sequence ambiguity is due to the gap in the *y*-fragments ladder between *y*_17_^+^ (the largest detected singly charged *y*-ion of sufficient abundance) and *y*_20_^2+^. The measured mass difference between these two fragments (after correction for the different protonation states) is 333.133 Da, which agrees with the DAF sequence suggested by the de novo sequencing algorithm well within 10 ppm (333.132 Da), but also permits another option, EGF, as well as permutations of both (other isobaric di- and tri-peptide residues can be ruled out, since their masses fall far outside of the acceptable 10 ppm range). Once again, careful inspection of the fragment ion spectrum reveals a plethora of abundant unassigned peaks that can be readily interpreted as internal fragments (labeled with gold and olive dots in **Figure 6B**). The most abundant group of internal fragments covers the PGLEDAFDLWGQ segment with a single-residue resolution (see the golden ladder on top of the peptide sequence in **Figure 6B**). This internal ladder allows the DAF/EGF ambiguity to be resolved with high confidence, confirming the sequence within the corresponding gap region.

The second group of internal fragments covers the APGLEDAFD segment, also at a single-residue resolution (although one of the fragments in this series, APGLED, is indistinguishable from its isomer belonging to the first group, PGLEDA). The ladder confirms that the residue preceding Pro^5^ is indeed alanine. Furthermore, two additional unique internal fragments AAPGLE and AAPGL allow the sequence to be further extended (with another alanine residue). This leaves only two possibilities, having either FA or AF as the N-terminal dipeptide residue (placing SM or MS – dipeptides isobaric to AF – at the N-terminus of the peptide would make the measured peptide ion mass fall outside of the 10 ppm range). Therefore, the only uncertainty in the peptide sequence shown in **Figure 6B** is the ordering of the first two amino acid residues. Lastly, one other issue with determining the HCDR3 structure is the absence of proteolytic peptides that span both the C-terminal segment of FR3 and the N-terminal segment of HCDR3. However, we notice that the length of the HCDR3 segment covered by the tryptic fragment shown in **Figure 6B** is thirteen residues, which is a canonical length for HCDR3,^37^ suggesting that the sequence shown in **Figures 3** and **6B** represents HCDR3 in its entirety.

### The Fab N-glycan resides within the FR3 segment of the heavy chain and is a result of a single mutation

Identification of all N-glycosylation sites within the VITT mAb was carried out by using ions corresponding to mono- and di-saccharides as glycopeptide reporters in the fragment ion spectra.^38^ Specifically, combinations of the following reporter ions were used: *m/z* 204.09 (HexNAc), *m/z* 274.10 (NeuAc-H_2_O), *m/z* 292.11 (NeuAc) and *m/z* 366.14 (Hex/HexNAc or HexNAc/Hex). In addition to identifying four glycopeptides representing the canonical N-glycosylation site within the C_H_2 domain of the antibody (see ***Supplementary Material*** for more information), application of this filter allowed another glycopeptide to be identified (**Figure 7**). A careful examination of its fragment ion spectrum (**Figure 7C**) revealed – in addition to the abundant low-*m/z* reporter ions – a characteristic glycan chain fragmentation pattern terminating at an ion peak with a measured mass of 978.52 Da, likely representing the intact “base” peptide with a completely stripped carbohydrate chain. De-*N*-glycosylation of the antibody prior to its proteolytic processing results in a complete disappearance of the glycopeptide signal (**Figure 7B**), while a novel fragment was detected whose mass corresponds to the deamidated version of the putative carbohydrate-free peptide at 978.52 Da (*vide supra*). The novel doubly charged ion (*m/z* 490.25) displays a prominent signal in the de-*N*-glycosylated sample (**Figure 7B**), while being absent in the digest of the intact antibody (**Figure 7A**). Collisional activation of this peptide ion generates a distinct fragmentation pattern (**Figure 7D**), from which its amino acid sequence can be readily deduced (LTLSDDTSK). Although there are two aspartic acid residues in this sequence (and each can be considered a result of deamidation due to the enzymatic *N*-glycan removal), it is the first of these two residues that would satisfy the *N*-glycosylation site criterion when converted back to asparagine, Asn-Xxx-Ser/Thr. Furthermore, aligning this peptide with the available gene sequences allows the matching sequence within the FR3 segment of IGHV2-26 to be identified (LTLSKDTSK), in which a single Lys/Asn mutation gives rise to a novel *N*-glycosylation site.

**Figure 7.**
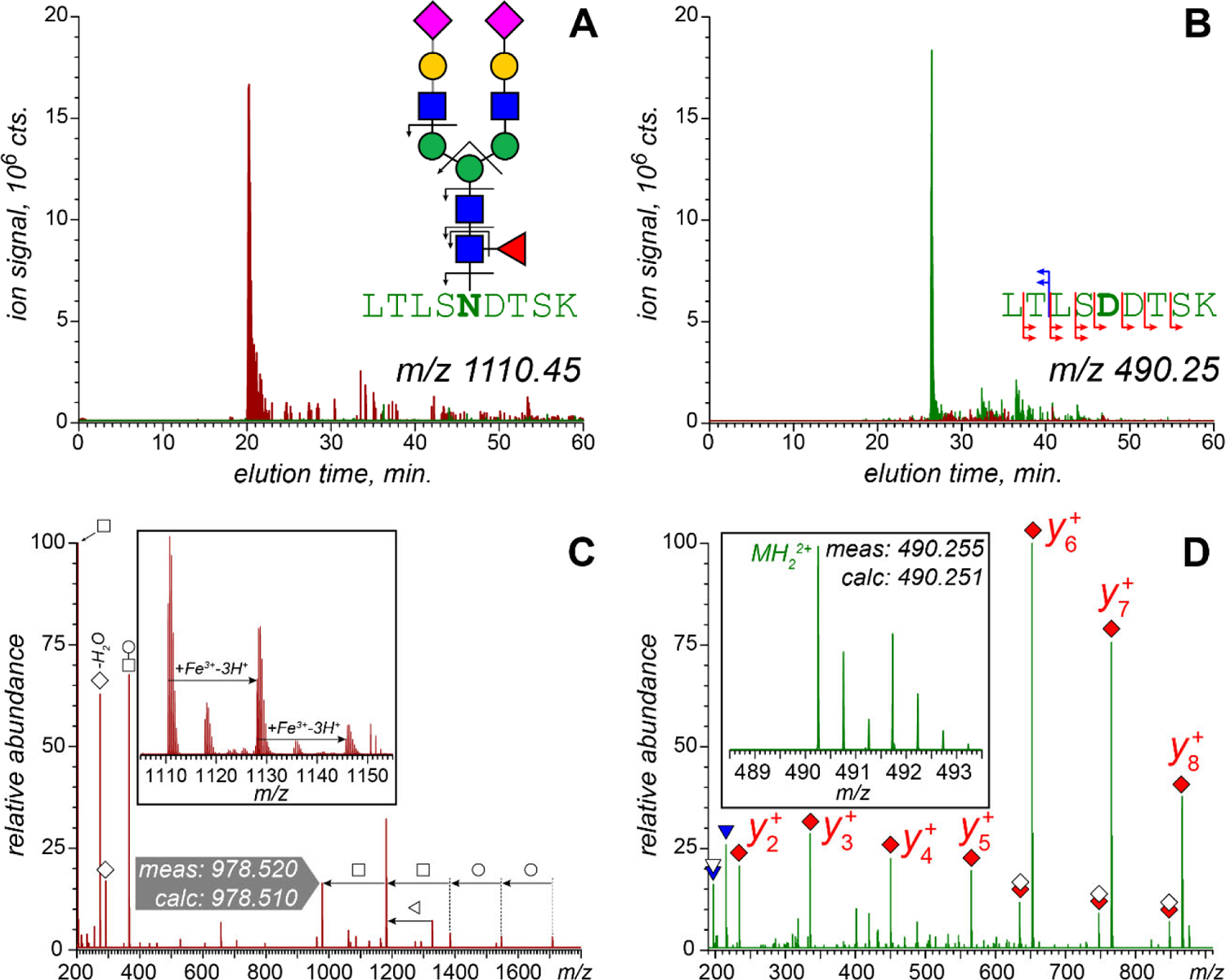
Localization of the N-glycosylation site within the Fab region of the VITT mAb by LC-MS/MS analysis of the tryptic fragment LTL(N*/D)TSK derived from the antibody before (maroon) and after (olive) PNGase F treatment. Panels **A** and **B** show TICs of the glycopeptide LTLN*TSK (*m/z* 1110.45) and its de-N-glycosylated form (*m/z* 490.25) in both samples, respectively. Panels **C** and **D** represent fragment ion spectra for these two peptides, and the framed insets show the molecular ions corresponding to these peptides in the MS1 spectra (which were used as the precursor ions for MS/MS measurements). The gray arrow in panel C identifies the ionic signal of the peptide from which the entire carbohydrate chain was stripped in the gas phase following collisional activation. The labeled low-mass ions (below *m/z* 400) in the same mass spectrum were used as “carbohydrate markers” when searching for glycopeptides in the entire LC-MS/MS data set.

### The polyclonal component of the VITT anti-PF4 antibodies is diverse and represented by all sub-classes of the heavy (IgG1 through IgG4) and light (both λ and κ) chains

The lower-abundance peptide fragments representing the polyclonal component of the antibody sample provide an opportunity to evaluate the diversity of antibodies outside of the monoclonal component. Multiple low-abundance fragments can be readily aligned with the IGKC gene,^35^ revealing the presence of the κ-type light chains. Furthermore, lower-abundance peptide fragments mapping to the hinge region in the heavy chain reveal the presence of all four subclasses of the latter (IgG1 through IgG4). While the quantitation of the fractions of these subclasses based on the ionic signal intensity is a dubious proposition, especially given the dramatic variation of the length and composition of the corresponding peptides, the minor subclass-specific variations within the peptides encompassing the canonical *N*-glycosylation site within the C_H_2 domain provide a viable opportunity vis-à-vis relative quantitation of the sub-classes. Indeed, the corresponding tryptic peptide in IgG2 (EEQ<u>F</u>N*ST<u>F</u>R) is distinct from both IgG1 (EEQ<u>Y</u>N*ST<u>Y</u>R), IgG3 (EEQ<u>Y</u>N*ST<u>F</u>R) and IgG4 (EEQ<u>F</u>N*ST<u>Y</u>R). Since the Phe/Tyr substitutions are not expected to result in a dramatic signal attenuation, the signal intensity can be used as a measure of the relative abundance of these sub-classes. This analysis (see ***Supplementary Material*** for more detail) confirms the prevalence of the IgG2 subclass in the antibody sample (over 80%), while the IgG1 content is only 10% (followed by a single-percentage point contribution from IgG3 and IgG4 combined). The sub-class distribution diagrams shown in the ***Supplementary Material*** are also noteworthy for allowing the limitations of this approach to the subclass quantitation to be defined. Indeed, while the 10:87:3 IgG1/IgG2/(IgG3+IgG4) ratio reported in the ***Supplementary Material*** is calculated based on the abundance of the fully protonated molecular ions (*MH*_4_^4+^), a notably lower fraction of IgG2 (73%) is obtained when the calculations are based on the intensity of *Fe*^3+^ adducts (*MHFe*^4+^). The *Fe*^3+^ adducts of carbohydrate ions have previously been reported by Klein and Zaia,^39^ and it appears that the presence of two additional hydroxyl groups within the glycopeptide derived from IgG1 (compared to IgG2) should increase the binding affinity by adding two potential ligands (hydroxyl groups), therefore decreasing the relative abundance of ions representing the IgG2-derived peptides.

Regardless of the procedure used to calculate the fractions of different IgG subclasses within the anti-PF4 antibodies extracted from a VITT patient’s plasma, IgG2 clearly is the most abundant class in this sample. While the low combined fraction of IgG3 and IgG4 in the antibody is not surprising (they typically account for ca. 10% of the total IgG content), the most abundant subclass in the human sera is IgG1,^40^ accounting for ca. 65% of the total. The dramatic distortion of the IgG2/(total IgG) balance observed in our work for the anti-PF4 IgGs extracted from a VITT patient’s blood is noteworthy, since the IgG2s are known to be involved primarily with recognizing polysaccharides,^41^ a class of biopolymers notably lacking within its presumed antigen, PF4. While the VITT mAb (which belongs to the IgG2 subclass, *vide supra*) clearly is one of the primary causes of this misbalance, it remains to be seen what the subclass distribution is among the polyclonal anti-PF4 antibodies, and whether it has any relevance vis-à-vis VITT etiology.

## Conclusions

Characterization of pathogenic anti-PF4 antibodies derived from a VITT patient’s blood using a variety of mass spectrometry-based approaches reveals several intriguing features. First, the subclass distribution is heavily biased towards IgG2, and in fact a significant proportion of the entire antibody ensemble is a monoclonal IgG2 molecule, VITT mAb. This is surprising, since the primary target of IgG2 is bacterial polysaccharides, and carbohydrate chains are notably absent within the VITT mAb’s presumed antigen (a small chemokine PF4). Carbohydrates (O-glycans) are present within the fiber proteins of Ad vectors,^42^ but their role in Ad immunogenicity remains obscure. Second, the amino acid sequences of both LCDR3 and HCDR3 (the two most variable segments in the light and heavy chains, respectively) show a significant similarity of the sequences reported earlier for an unrelated set of VITT patients.^12^ This similarity is particularly noteworthy for HCDR3, since it is produced by recombination of three genes, V_H_, D and J, and is the most variable antibody segment even in the absence of somatic hypermutations. Lastly, VITT mAb has a fully mature biantennary *N*-glycan within its V_H_ domain (FR3). While the presence of *N*-glycans in IgG molecules outside of the C_H_2 domain is currently viewed as uncommon,^27^ recent reports suggest that this structural feature is shared by several autoimmune disorders.^43^ In fact, it may serve as a molecular trigger that activates non-canonical clone selection pathways and allows it to evade elimination by the elaborate self/not-self control machinery,^29^ thus leading to the onset of an autoimmune disorder.

## Authors contributions

SNN and SHL planned and carried out experimental work and data interpretation on VITT mAb large fragment characterization, as well as antibody sequencing and N-glycan localization; DGI planned and carried out intact-mass measurements and participated in sequencing data interpretation; NI and IN provided the antibody sample; IAK designed the study, participated in data interpretation and wrote the manuscript. All authors participated in editing the manuscript and gave their consent to submitting its final version for peer-review and publication.

## Acknowledgments

This work was supported by a grant R01 GM112666 from the National Institute of Health. All MS measurements were carried out in the Mass Spectrometry Core facility at UMass-Amherst (RRID:SCR_019063).

## Supplementary Material for

**Figure S1.**
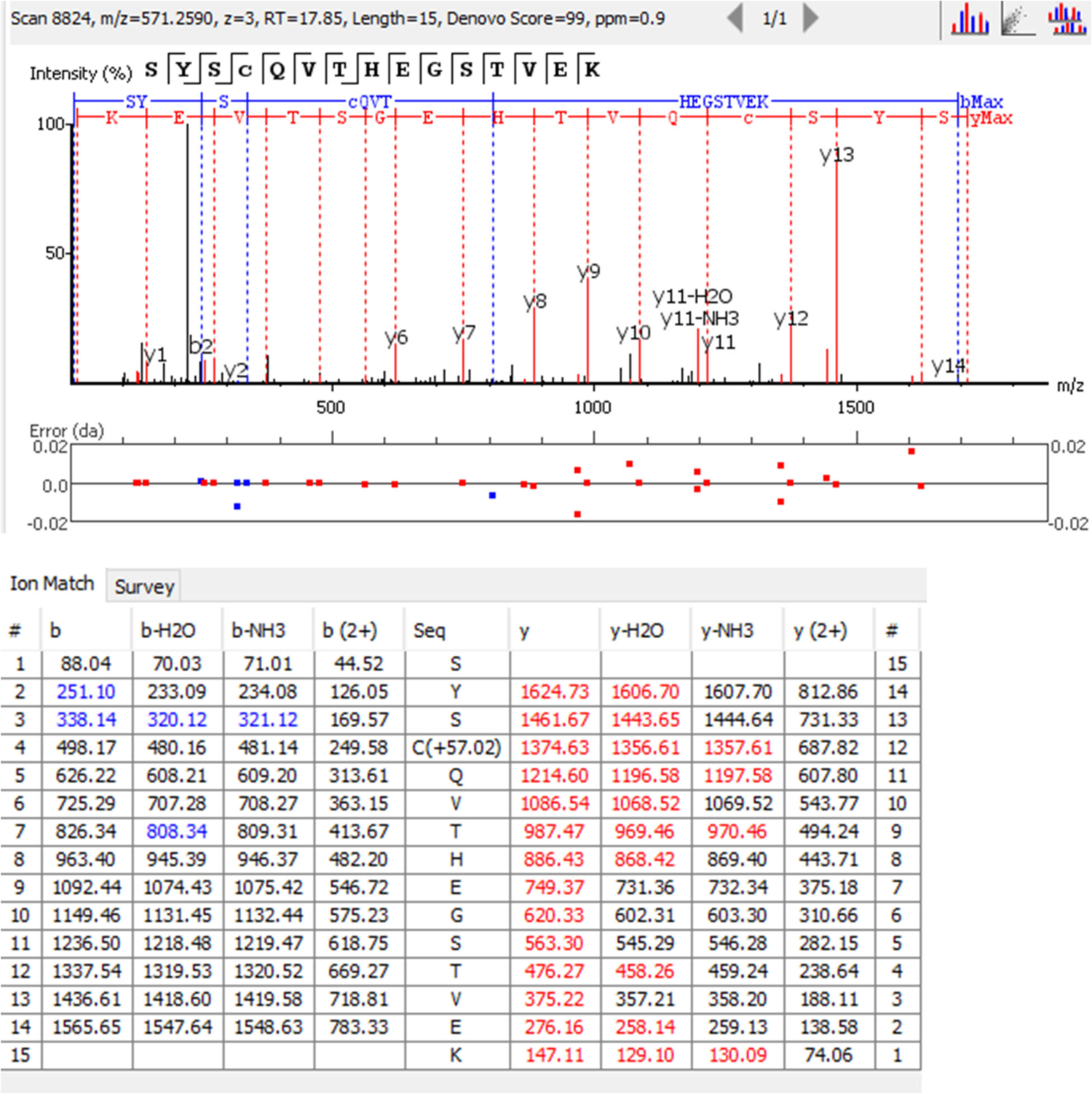
An example of LC-MS/MS analysis of a peptide derived from the C_L_ domain of VITT mAb confirming the light chain isotype assignment λ (tryptic fragment Ser^L191^→Lys^L205^).

**Figure S2.**
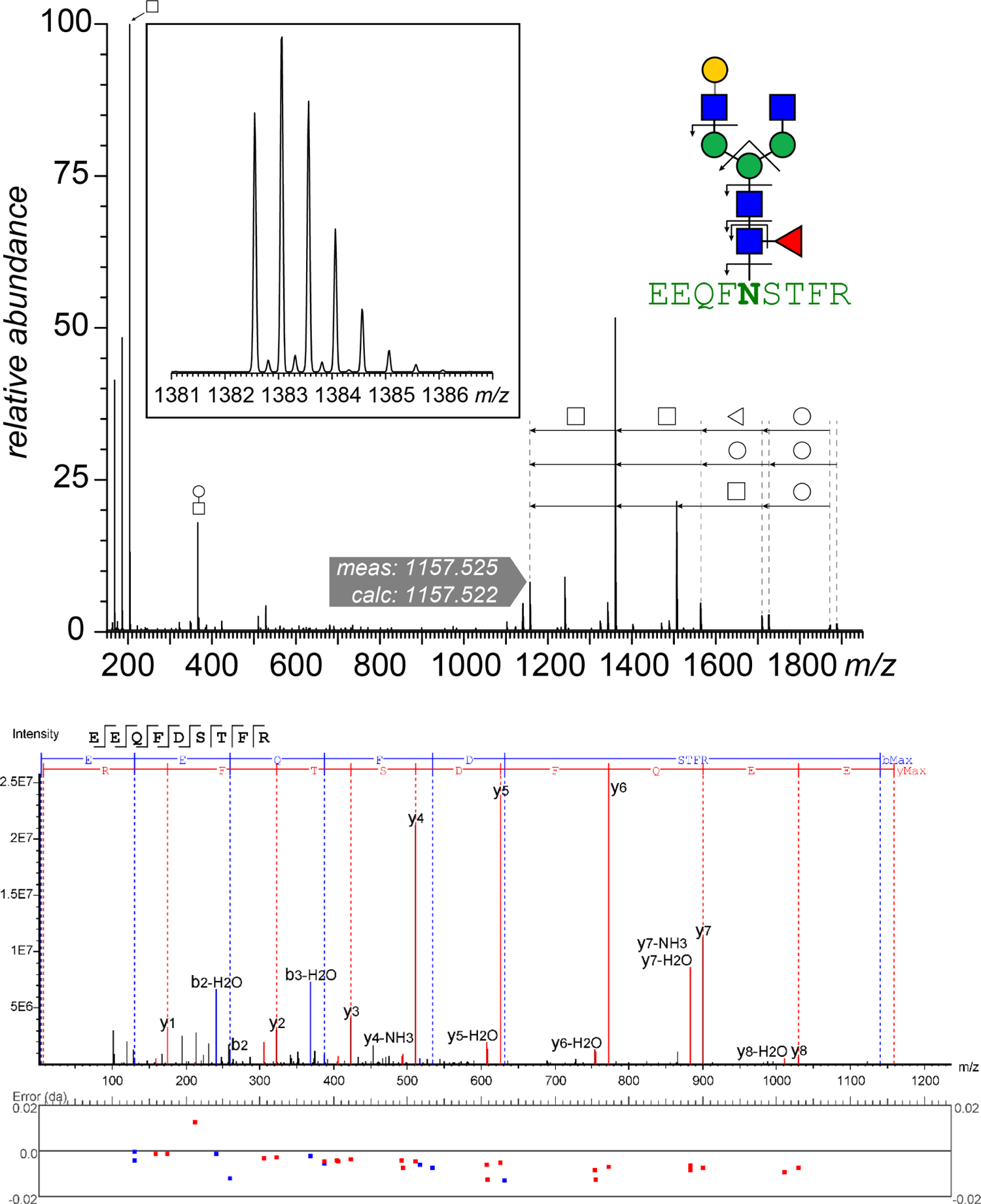
LC-MS/MS analysis of a glycopeptide derived from the C_H_2 domain of VITT mAb before (top) and after de-N-glycosylation (bottom).

**Figure S3.**
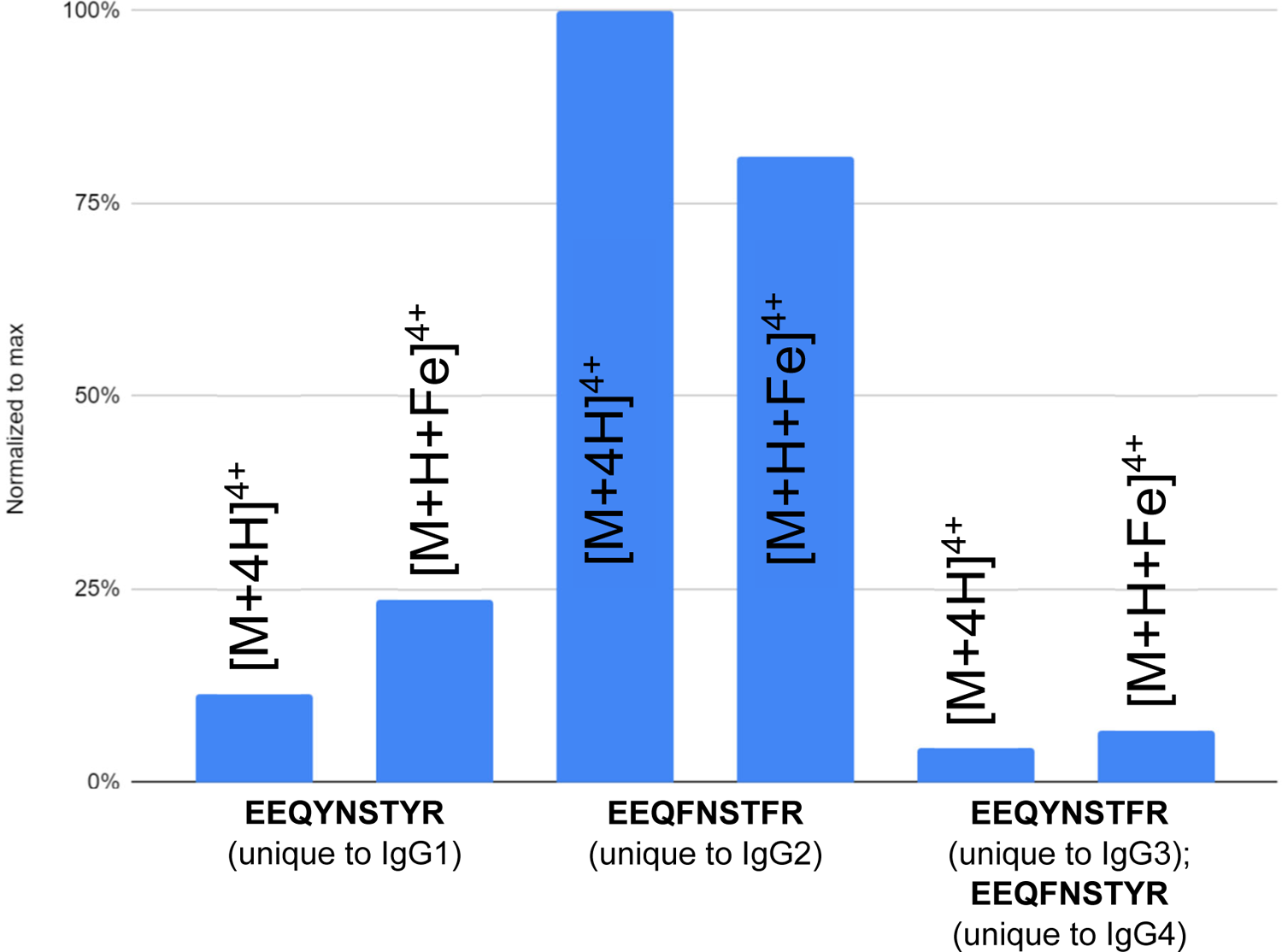
Quantitation of the anti-PF4 IgG subclasses in the clinical sample based on the abundance of unique glycopeptide ions representing the canonical *N*-glycosylation site in the C_H_2 domain.

## References

1. Chan, B. T.; Bobos, P.; Odutayo, A.; Pai, M. Meta-Analysis of Risk of Vaccine-Induced Immune Thrombotic Thrombocytopenia Following ChAdOx1-S Recombinant Vaccine. medRxiv 2021, 2021.2005.2004.21256613. DOI: 10.1101/2021.05.04.21256613%JmedRxiv.

2. Akinshina, S.; Bitsadze, V.; Khizroeva, J. K.; Grigorieva, K.; Slukhanchuk, E.; Tretyakova, M.; Tsibizova, V.; Elalamy, I.; Gris, J.-C.; Brenner, B. J. O., Gynecology;, et al. Vaccine-induced immune thrombotic thrombocytopenia: definition, risks with different vaccines, and regulatory responses. *Akusherstvo, Ginekologia i Reprodukcia (Obstetrics*, Gynecology and Reproduction) 2021, 15 (5), 562-575. DOI: https://doi.org/10.17749/2313-7347/ob.gyn.rep.2021.257. Jevtic, S. D.; Arnold, D. M.; Modi, D.; Ivetic, N.; Bissola, A. L.; Nazy, I. Vaccine-induced immune thrombotic thrombocytopenia: Updates in pathobiology and diagnosis. *Front. Cardiovasc. Med.* 2022, *9*, 1040196. DOI: 10.3389/fcvm.2022.1040196 From NLM.

3. Herrera-Comoglio, R.; Lane, S. Vaccine-Induced Immune Thrombocytopenia and Thrombosis after the Sputnik V Vaccine. N. Engl. J. Med. 2022, 387 (15), 1431-1432. DOI: 10.1056/NEJMc2210813 From NLM.

4. Schultz, N. H.; Sørvoll, I. H.; Michelsen, A. E.; Munthe, L. A.; Lund-Johansen, F.; Ahlen, M. T.; Wiedmann, M.; Aamodt, A. H.; Skattør, T. H.; Tjønnfjord, G. E.;, et al. Thrombosis and Thrombocytopenia after ChAdOx1 nCoV-19 Vaccination. N. Engl. J. Med. 2021, 384 (22), 2124-2130. DOI: 10.1056/NEJMoa2104882 From NLM.

5. Zidan, A.; Noureldin, A.; Kumar, S. A.; Elsebaie, A.; Othman, M. COVID-19 Vaccine-Associated Immune Thrombosis and Thrombocytopenia (VITT): Diagnostic Discrepancies and Global Implications. Semin. Thromb. Hemost. 2023, 49 (1), 9-14. DOI: 10.1055/s-0042-1759684 From NLM. Adu, P.; Poopola, T.; Medvedev, O. N.; Collings, S.; Mbinta, J.; Aspin, C.; Simpson, C. R. Implications for COVID-19 vaccine uptake: A systematic review. *J. Infect. Public Health* 2023, *16* (3), 441-466. DOI: 10.1016/j.jiph.2023.01.020 From NLM. Abad, N.; Messinger, S. D.; Huang, Q.; Hendrich, M. A.; Johanson, N.; Fisun, H.; Lewis, Z.; Wilhelm, E.; Baack, B.; Bonner, K. E.; et al. A qualitative study of behavioral and social drivers of COVID-19 vaccine confidence and uptake among unvaccinated Americans in the US April-May 2021. *PloS one* 2023, *18* (2), e0281497. DOI: 10.1371/journal.pone.0281497 From NLM.

6. Coughlan, L.; Kremer, E. J.; Shayakhmetov, D. M. Adenovirus-based vaccines-a platform for pandemic preparedness against emerging viral pathogens. Mol. Ther. 2022, 30 (5), 1822-1849. DOI: 10.1016/j.ymthe.2022.01.034 From NLM.

7. Salih, F.; Schonborn, L.; Endres, M.; Greinacher, A. Immunvermittelte Sinus- und Hirnvenenthrombosen: VITT und prä-VITT als Modellerkrankung [Cerebral Venous and Sinus Thrombosis with Immunological Pathogenesis: VITT and Pre-VITT]. Aktuelle Rheumatol. 2022, 47, 490-501, Article; Early Access. DOI: 10.1055/a-1936-3123. Lippi, G.; Favaloro, E. J. Cerebral Venous Thrombosis Developing after COVID-19 Vaccination: VITT, VATT, TTS, and More. *Semin. Thromb. Hemost.* 2022, *48* (1), 8-14. DOI: 10.1055/s-0041-1736168 From NLM.

8. Chen, L.; Pavord, S. Clinical picture of VITT. Semin. Hematol. 2022, 59 (2), 76-79. DOI: 10.1053/j.seminhematol.2022.02.001 From NLM.

9. Scully, M.; Singh, D.; Lown, R.; Poles, A.; Solomon, T.; Levi, M.; Goldblatt, D.; Kotoucek, P.; Thomas, W.; Lester, W. Pathologic Antibodies to Platelet Factor 4 after ChAdOx1 nCoV-19 Vaccination. N. Engl. J. Med. 2021, 384 (23), 2202-2211. DOI: 10.1056/NEJMoa2105385 From NLM.

10. Huynh, A.; Kelton, J. G.; Arnold, D. M.; Daka, M.; Nazy, I. Antibody epitopes in vaccine-induced immune thrombotic thrombocytopenia. Nature 2021, 596, 565-569. DOI: 10.1038/s41586-021-03744-4 From NLM.

11. Kanack, A. J.; Bayas, A.; George, G.; Abou-Ismail, M. Y.; Singh, B.; Kohlhagen, M. C.; Splinter, N. P.; Christ, M.; Naumann, M.; Moser, K. A.;, et al. Monoclonal and oligoclonal anti-platelet factor 4 antibodies mediate VITT. Blood 2022, 140 (1), 73-77. DOI: 10.1182/blood.2021014588 From NLM.

12. Wang, J. J.; Armour, B.; Chataway, T.; Troelnikov, A.; Colella, A.; Yacoub, O.; Hockley, S.; Tan, C. W.; Gordon, T. P. Vaccine-induced immune thrombotic thrombocytopenia is mediated by a stereotyped clonotypic antibody. Blood 2022, 140 (15), 1738-1742. DOI: 10.1182/blood.2022016474 From NLM.

13. van de Bovenkamp, F. S.; Hafkenscheid, L.; Rispens, T.; Rombouts, Y. The Emerging Importance of IgG Fab Glycosylation in Immunity. J. Immunol. 2016, 196 (4), 1435-1441. DOI: 10.4049/jimmunol.1502136 From NLM.

14. Goldstein, L. D.; Chen, Y. J.; Wu, J.; Chaudhuri, S.; Hsiao, Y. C.; Schneider, K.; Hoi, K. H.; Lin, Z.; Guerrero, S.; Jaiswal, B. S.;, et al. Massively parallel single-cell B-cell receptor sequencing enables rapid discovery of diverse antigen-reactive antibodies. *Commun*. Biol. 2019, 2, 304. DOI: 10.1038/s42003-019-0551-y From NLM.

15. Guthals, A.; Gan, Y.; Murray, L.; Chen, Y.; Stinson, J.; Nakamura, G.; Lill, J. R.; Sandoval, W.; Bandeira, N. De Novo MS/MS Sequencing of Native Human Antibodies. J. Proteome Res. 2017, 16 (1), 45-54. DOI: 10.1021/acs.jproteome.6b00608. Peng, W.; Pronker, M. F.; Snijder, J. Mass Spectrometry-Based De Novo Sequencing of Monoclonal Antibodies Using Multiple Proteases and a Dual Fragmentation Scheme. *J. Proteome Res.* 2021, *20* (7), 3559-3566. DOI: 10.1021/acs.jproteome.1c00169 From NLM. Melani, R. D.; Des Soye, B. J.; Kafader, J. O.; Forte, E.; Hollas, M.; Blagojevic, V.; Negrão, F.; McGee, J. P.; Drown, B.; Lloyd-Jones, C.; et al. Next-Generation Serology by Mass Spectrometry: Readout of the SARS-CoV-2 Antibody Repertoire. *J. Proteome Res.* 2022, *21* (1), 274-288. DOI: 10.1021/acs.jproteome.1c00882 From NLM. de Graaf, S. C.; Hoek, M.; Tamara, S.; Heck, A. J. R. A perspective toward mass spectrometry-based de novo sequencing of endogenous antibodies. *mAbs* 2022, *14* (1), 2079449. DOI: 10.1080/19420862.2022.2079449.

16. Beslic, D.; Tscheuschner, G.; Renard, B. Y.; Weller, M. G.; Muth, T. Comprehensive evaluation of peptide de novo sequencing tools for monoclonal antibody assembly. Brief. Bioinform. 2023, 24 (1), 1-12. DOI: 10.1093/bib/bbac542 From NLM.

17. Huynh, A.; Arnold, D. M.; Smith, J. W.; Elliott, T. D.; Ivetic, N.; Kelton, J. G.; Nazy, I. The role of fluid-phase immune complexes in the pathogenesis of heparin-induced thrombocytopenia. Thromb. Res. 2020, 194, 135-141. DOI: 10.1016/j.thromres.2020.06.012 From NLM.

18. Abzalimov, R. R.; Kaltashov, I. A. Electrospray ionization mass spectrometry of highly heterogeneous protein systems: Protein ion charge state assignment via incomplete charge reduction. Anal. Chem. 2010, 82 (18), 7523-7526, Article. DOI: 10.1021/ac101848z.

19. Marty, M. T.; Baldwin, A. J.; Marklund, E. G.; Hochberg, G. K.; Benesch, J. L.; Robinson, C. V. Bayesian deconvolution of mass and ion mobility spectra: from binary interactions to polydisperse ensembles. Anal. Chem. 2015, 87 (8), 4370-4376. DOI: 10.1021/acs.analchem.5b00140 From NLM.

20. Bobst, C. E.; Sperry, J.; Friese, O. V.; Kaltashov, I. A. Simultaneous Evaluation of a Vaccine Component Microheterogeneity and Conformational Integrity Using Native Mass Spectrometry and Limited Charge Reduction. J. Am. Soc. Mass Spectrom. 2021, 32 (7), 1631-1637. DOI: 10.1021/jasms.1c00091. Kafader, J. O.; Melani, R. D.; Schachner, L. F.; Ives, A. N.; Patrie, S. M.; Kelleher, N. L.; Compton, P. D. Native vs Denatured: An in Depth Investigation of Charge State and Isotope Distributions. *J. Am. Soc. Mass Spectrom.* 2020, *31* (3), 574-581. DOI: 10.1021/jasms.9b00040.

21. Kaltashov, I. A.; Ivanov, D. G.; Yang, Y. Mass spectrometry-based methods to characterize highly heterogeneous biopharmaceuticals, vaccines, and nonbiological complex drugs at the intact-mass level. Mass Spectrom. Rev. 2022, e21829. DOI: 10.1002/mas.21829 From NLM.

22. Yang, W.; Ivanov, D. G.; Kaltashov, I. A. Extending the capabilities of intact-mass analyses to monoclonal immunoglobulins of the E-isotype (IgE). mAbs 2022, 14 (1), 2103906. DOI: 10.1080/19420862.2022.2103906 From NLM.

23. Huynh, A.; Arnold, D. M.; Kelton, J. G.; Smith, J. W.; Horsewood, P.; Clare, R.; Guarne, A.; Nazy, I. Characterization of platelet factor 4 amino acids that bind pathogenic antibodies in heparin-induced thrombocytopenia. J. Thromb. Haemost. 2019, 17 (2), 389-399. DOI: 10.1111/jth.14369 From NLM.

24. Pawlowski, J. W.; Bajardi-Taccioli, A.; Houde, D.; Feschenko, M.; Carlage, T.; Kaltashov, I. A. Influence of glycan modification on IgG1 biochemical and biophysical properties. J. Pharm. Biomed. Anal. 2018, 151, 133-144. DOI: https://doi.org/10.1016/j.jpba.2017.12.061.

25. Barnidge, D. R.; Dasari, S.; Ramirez-Alvarado, M.; Fontan, A.; Willrich, M. A.; Tschumper, R. C.; Jelinek, D. F.; Snyder, M. R.; Dispenzieri, A.; Katzmann, J. A.;, et al. Phenotyping polyclonal kappa and lambda light chain molecular mass distributions in patient serum using mass spectrometry. J. Proteome Res. 2014, 13 (11), 5198-5205. DOI: 10.1021/pr5005967 From NLM.

26. Damelang, T.; Rogerson, S. J.; Kent, S. J.; Chung, A. W. Role of IgG3 in Infectious Diseases. Trends Immunol. 2019, 40 (3), 197-211. DOI: 10.1016/j.it.2019.01.005 From NLM.

27. Maverakis, E.; Kim, K.; Shimoda, M.; Gershwin, M. E.; Patel, F.; Wilken, R.; Raychaudhuri, S.; Ruhaak, L. R.; Lebrilla, C. B. Glycans in the immune system and The Altered Glycan Theory of Autoimmunity: a critical review. J. Autoimmun. 2015, 57, 1-13. DOI: 10.1016/j.jaut.2014.12.002 From NLM.

28. Vergroesen, R. D.; Slot, L. M.; Hafkenscheid, L.; Koning, M. T.; van der Voort, E. I. H.; Grooff, C. A.; Zervakis, G.; Veelken, H.; Huizinga, T. W. J.; Rispens, T.;, et al. B-cell receptor sequencing of anti-citrullinated protein antibody (ACPA) IgG-expressing B cells indicates a selective advantage for the introduction of N-glycosylation sites during somatic hypermutation. Ann. Rheum. Dis. 2018, 77 (6), 956-958. DOI: 10.1136/annrheumdis-2017-212052 From NLM. Kissel, T.; Ge, C.; Hafkenscheid, L.; Kwekkeboom, J. C.; Slot, L. M.; Cavallari, M.; He, Y.; van Schie, K. A.; Vergroesen, R. D.; Kampstra, A. S. B.; et al. Surface Ig variable domain glycosylation affects autoantigen binding and acts as threshold for human autoreactive B cell activation. *Sci. Adv.* 2022, *8* (6), eabm1759. DOI: 10.1126/sciadv.abm1759 From NLM. Koers, J.; Sciarrillo, R.; Derksen, N. I. L.; Vletter, E. M.; Fillié-Grijpma, Y. E.; Raveling-Eelsing, E.; Graça, N. A. G.; Leijser, T.; Pas, H. H.; Laura van Nijen-Vos, L.; et al. Differences in IgG autoantibody Fab glycosylation across autoimmune diseases. *J. Allergy Clin. Immunol.* 2023, *in press, doi: 10.1016/j.jaci.2022.10.035*. DOI: 10.1016/j.jaci.2022.10.035 From NLM.

29. Visser, A.; Hamza, N.; Kroese, F. G. M.; Bos, N. A. Acquiring new N-glycosylation sites in variable regions of immunoglobulin genes by somatic hypermutation is a common feature of autoimmune diseases. Ann. Rheum. Dis. 2018, 77 (10), e69. DOI: 10.1136/annrheumdis-2017-212568 From NLM.

30. Gu, J.; Lei, Y.; Huang, Y.; Zhao, Y.; Li, J.; Huang, T.; Zhang, J.; Wang, J.; Deng, X.; Chen, Z.;, et al. Fab fragment glycosylated IgG may play a central role in placental immune evasion. Hum. Reprod. 2015, 30 (2), 380-391. DOI: 10.1093/humrep/deu323 From NLM.

31. Courtois, F.; Agrawal, N. J.; Lauer, T. M.; Trout, B. L. Rational design of therapeutic mAbs against aggregation through protein engineering and incorporation of glycosylation motifs applied to bevacizumab. mAbs 2016, 8 (1), 99-112. DOI: 10.1080/19420862.2015.1112477 From NLM. van de Bovenkamp, F. S.; Derksen, N. I. L.; van Breemen, M. J.; de Taeye, S. W.; Ooijevaar-de Heer, P.; Sanders, R. W.; Rispens, T. Variable Domain N-Linked Glycans Acquired During Antigen-Specific Immune Responses Can Contribute to Immunoglobulin G Antibody Stability. *Front. Immunol.* 2018, *9*, 740. DOI: 10.3389/fimmu.2018.00740 From NLM.

32. Bruhns, P.; Iannascoli, B.; England, P.; Mancardi, D. A.; Fernandez, N.; Jorieux, S.; Daëron, M. Specificity and affinity of human Fcgamma receptors and their polymorphic variants for human IgG subclasses. Blood 2009, 113 (16), 3716-3725. DOI: 10.1182/blood-2008-09-179754 From NLM.

33. Scherer, H. U.; van der Woude, D.; Toes, R. E. M. From risk to chronicity: evolution of autoreactive B cell and antibody responses in rheumatoid arthritis. Nat. Rev. Rheumatol. 2022, 18 (7), 371-383. DOI: 10.1038/s41584-022-00786-4 From NLM.

34. Tran, N. H.; Qiao, R.; Xin, L.; Chen, X.; Liu, C.; Zhang, X.; Shan, B.; Ghodsi, A.; Li, M. Deep learning enables de novo peptide sequencing from data-independent-acquisition mass spectrometry. Nat. Methods 2019, 16 (1), 63-66. DOI: 10.1038/s41592-018-0260-3 From NLM.

35. Lefranc, M. P.; Ehrenmann, F.; Kossida, S.; Giudicelli, V.; Duroux, P. Use of IMGT(®) Databases and Tools for Antibody Engineering and Humanization. Methods Mol. Biol. 2018, 1827, 35-69. DOI: 10.1007/978-1-4939-8648-4_3 From NLM.

36. Murphy, K.; Travers, P.; Walport, M.; Janeway, C. Janeway’s immunobiology; Garland Science, 2012.

37. Chiu, M. L.; Goulet, D. R.; Teplyakov, A.; Gilliland, G. L. Antibody Structure and Function: The Basis for Engineering Therapeutics. Antibodies 2019, 8 (4), 55. DOI: 10.3390/antib8040055 From NLM.

38. Kaltashov, I. A.; Wang, S.; Wang, G. Mass Spectrometry in Biopharmaceutical Analysis; De Gruyter, 2021. DOI: 10.1515/9783110546187.

39. Klein, J.; Zaia, J. Relative Retention Time Estimation Improves N-Glycopeptide Identifications by LC-MS/MS. J. Proteome Res. 2020, 19 (5), 2113-2121. DOI: 10.1021/acs.jproteome.0c00051 From NLM.

40. Papadea, C.; Check, I. J. Human immunoglobulin G and immunoglobulin G subclasses: biochemical, genetic, and clinical aspects. Crit. Rev. Clin. Lab. Sci. 1989, 27 (1), 27-58. DOI: 10.3109/10408368909106589 From NLM.

41. Vidarsson, G.; Dekkers, G.; Rispens, T. IgG subclasses and allotypes: from structure to effector functions. Front. Immunol. 2014, 5, 520. DOI: 10.3389/fimmu.2014.00520 From NLM.

42. Cauet, G.; Strub, J.-M.; Leize, E.; Wagner, E.; Dorsselaer, A. V.; Lusky, M. Identification of the Glycosylation Site of the Adenovirus Type 5 Fiber Protein. Biochemistry 2005, 44 (14), 5453-5460. DOI: 10.1021/bi047702b.

43. Vletter, E. M.; Koning, M. T.; Scherer, H. U.; Veelken, H.; Toes, R. E. M. A Comparison of Immunoglobulin Variable Region N-Linked Glycosylation in Healthy Donors, Autoimmune Disease and Lymphoma. Front. Immunol. 2020, 11, 241. DOI: 10.3389/fimmu.2020.00241 From NLM.

